# Learning to act by integrating mental simulations and physical experiments

**DOI:** 10.1101/321497

**Authors:** Ishita Dasgupta, Kevin A. Smith, Eric Schulz, Joshua B. Tenenbaum, Samuel J. Gershman

**Author notes:** These authors contributed equally to the work (Ishita Dasgupta, Kevin A. Smith).

## Abstract

People can learn about the effects of their actions either by performing physical experiments or by running mental simulations. Physical experiments are reliable but risky; mental simulations are unreliable but safe. We investigate how people negotiate the balance between these strategies. Participants attempted to shoot a ball at a target, and could pay to take practice shots (physical experiments). They could also simply think (run mental simulations), but were incentivized to act quickly by paying for time. We demonstrate that the amount of thinking time and physical experiments is sensitive to trial characteristics in a way that is consistent with a model that integrates information across simulation and experimentation and decides online when to perform each.

## Introduction

Craik (1943) famously proposed that an advanced organism might hold “a ‘small-scale model’ of external reality and of its own possible actions in its head… to try out various alternatives, conclude which is the best of them… [and] to react in a much fuller, safer, and more competent manner.” These simulatable models allow us to gain information about the effect of our actions without needing to incur the cost of performing those actions. Simulation using mental models has explained how we perform not just spatial (Kosslyn et al., 1978; Shepard & Metzler, 1971) and physical (Battaglia et al., 2013; Hegarty, 2004) reasoning, but has also been suggested to underlie language understanding (Bergen, 2012) and theory of mind (Gallese & Goldman, 1998).

While simulations are cheaper, safer, and faster than acting on the world, they are not perfect substitutes. Because any mental model only approximates the world, simulation provides uncertain predictions of the effects of our actions (Smith & Vul, 2013). Furthermore, simulations are less costly than actions but are not costless – they require mental effort and time to perform (Vul et al., 2014). Thus, when we want to learn about the world, we need to decide how to combine relatively cheap and imprecise simulations with more costly and accurate physical experiments. To do this rationally requires integrating information from both simulation and action and assessing their expected benefits and costs (Gershman et al., 2015; Lieder & Griffiths, 2017).

We study this problem in a physical action planning task, which serves as a suitable test bed because we have models of uncertainty in people’s physical predictions (Smith & Vul, 2013), and because there is evidence that people make rational trade-offs about how much simulation to use when physical experiments are not possible (Hamrick et al., 2015). We use a task in which participants are asked to aim a ball at a goal, but can think about it or take practice shots beforehand (Fig. 1). We find that people modulate the number of experiments and time spent thinking based on trial characteristics that cannot be explained solely by simple measures of difficulty.

We then propose a model of how people combine simulation and experimentation. This model is based on the theory that just as we can learn mappings between actions and outcomes by real-world observations, we can gain information about this mapping using simulations in place of observations (Gershman et al., 2017; Sutton, 1991). This model uses a type of Bayesian optimization – Gaussian process entropy optimization (Hernández-Lobato et al., 2014) – that integrates different sources of information and decides between performing noisy but cheap simulations and running more expensive but near-deterministic experiments. It captures aspects of how many experiments people perform, how much time they spend thinking, as well as their ultimate action choices.

**Figure 1:**
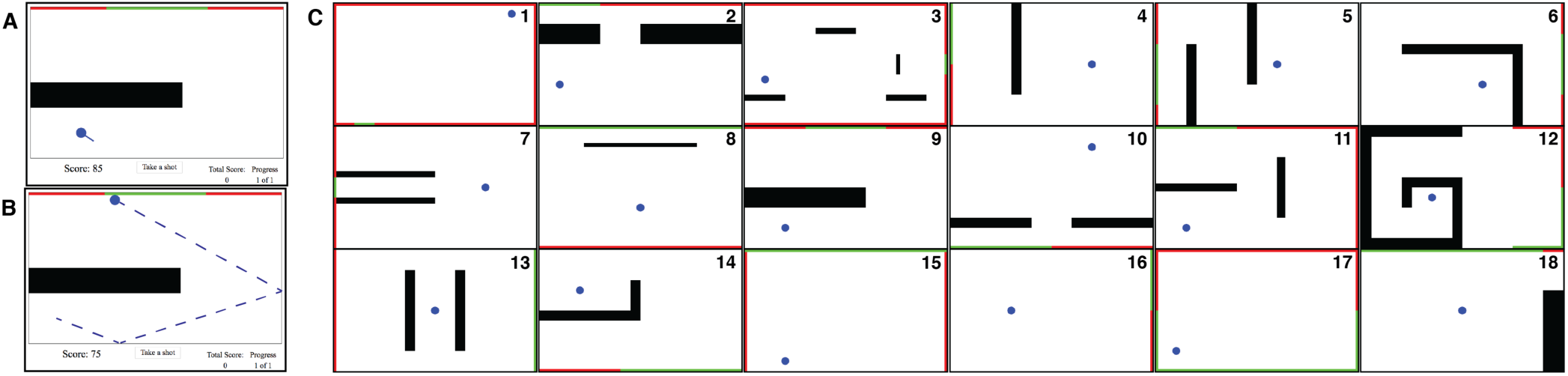
Diagram of the experimental trials. **A**: Participants observe a table and are asked to shoot the ball into the green goal without hitting the red areas. Participants could aim the ball and take practice shots at a cost to the score. **B**: The trajectory and outcome of shots were shown. Once participants had finished practicing and thinking, they could click the ‘Take a shot’ button and would have one chance to score. **C**: The 18 trials used in the experiment, ordered from least to most likely to succeed by chance.

## Experiment

We developed a physical reasoning task in which participants choose an angle to launch a ball at a target. On each trial (see Fig. 1), they could choose to perform physical experiments as well as think about how to shoot the ball. Once they decided they were ready, they had to perform an action in the exact same scenario and earned points based on the success of their action as well as on how much time they had spent thinking about the task and how many experiments they had run.

**Participants.** We recruited 60 participants from Amazon Mechanical Turk using the psiTurk framework (Gureckis et al., 2016). Each participant was paid a participation fee of $2.50 and randomly assigned to the *cheap experiments* condition or *costly experiments* condition.

**Procedure.** On each trial, participants observed a rectangular table of 1000 × 600 pixels on their screen. On this table was a single blue ball, any number of black walls, and on the edge of the table were green and red colored patches (see Fig 1). Participants were instructed that their task was to determine the angle to launch the ball so that it would touch the green goal before touching the red area or traveling too far a distance.

Each trial was divided into two phases: information gathering and the final action phase. In the first phase, participants started with 100 points. They were instructed that they could think or take practice shots, but both would cost points. The score would decay at a rate of 6 points/second, and each practice shot would incur a cost depending on the experimental condition: 10 points in the cheap condition and 20 points in the costly condition.

Participants could take a practice shot by positioning their mouse within 250px of the center of the ball and clicking. If the mouse was within this range, a straight line would extend from the ball indicating the direction of the shot. Once a practice shot was clicked, the ball would travel in that direction, bouncing off of any walls or table edges until it either reached the green or red areas, or had traveled a distance of 3000px (which would take ∼ 1.5s, though could vary based on the participant’s computer speed). During a practice shot, the score decay was paused.

When participants believed they knew how to shoot the ball, they could click a button below the table to move to the final phase. As soon as this button was clicked, the score would stop decaying, and a yellow border would appear around the table to indicate the final action phase. Participants would have three seconds to take their final shot, which could be performed in the same way as a practice shot. If this final shot hit the green goal, the participant would earn a score equal to the number of points remaining from the information gathering phase. If the final shot hit the red area or timed out, the participant would lose 10 points from their total score. If the participant did not make a shot within the three second limit, they would also lose 10 points. Afterwards, the participant would be notified of the outcome and move on to the next trial.

If the score decayed to zero during the information gathering phase, the final shot phase would immediately begin (including the yellow border notification). Participants would then earn no points for a successful shot, but could lose points for a miss or failure to act.

The total score was displayed in the lower right of the screen as a motivation for participants, but did not affect compensation.

**Materials.** Participants performed a total of 25 trials. The first seven trials were always presented in the same order and were designed as introductory trials to familiarize participants with the interface; these trials were not included in any analyses. The remaining 18 trials (see Fig. 1, right) were presented in a randomized order for each participant. These trials were designed by hand to provide a range of difficulties for an agent that took shots at random angles: six trials would be solved by fewer than 10% of random shots, six trials would be solved by between 10–30% of random shots, and six trials would be solved by greater than 30% of random shots. This probability of succeeding by guessing is used as a rough proxy for difficulty, and to assess whether participants were adapting their practice experiments or actual shots to new information or simply guessing. Trials were labeled by number in increasing probability of guessing.

**Data exclusions.** We removed data from three participants for whom we did not record actions for all 18 trials, and from seven additional participants who ‘timed out’ in either the information seeking or final action phase on more than half of the trials (indicating low attention). This left 27 participants in the cheap condition and 23 participants in the costly condition. We further excluded from analysis any trials in which the participants timed out only for the final shot (5.7%) and trials in which the participant minimized or browsed away from the experimental window (0.1%).

### Behavioral results

Overall, participants performed well in this task and hit the target on their final shot more frequently than would be expected under chance performance (61% accuracy versus 25% accuracy at chance; *t*(17) = 5.01, *p* < .001, *d* = 2.43).

There was no good evidence that participants performed fewer practice shots on average between the two experimental conditions (cheap: 0.68, costly: 0.48, *t*(47.3)= 1.08, *p* = .29, *d* = 0.31), nor was there evidence that the average time spent thinking differed by condition (cheap: 2.5*s*, costly: 2.4*s*, *t*(46.2) = 0.26, *p* = .79, *d* = 0.08). Participants were also roughly equally accurate across the two conditions (cheap: 63%, costly: 60%, *t*(40.0) = 0.82, *p* = .42, *d* = 0.26). Because we found no difference in behavior by cost condition, we ignore this manipulation for all future analyses.

Across individuals, simulation and experimentation were tightly linked: the average time participants spent thinking on a trial (excluding time watching practice shots) was correlated with the number of practice shots they took (*r*(48) = 0.76, *p* < .001).

At the trial level, participants were sensitive to the gross difficulty: as the probability of shooting the ball in the goal by chance increased they were more accurate (averaged over trials, *r*(16) = 0.72, *p* < .001), performed fewer experimental shots (*r* (16) = −0.52, *p* = .027, see Fig. 2, left), and took less time to think (*r*(16) = −0.65, *p* = .004, see Fig. 2, right).

**Figure 2:**
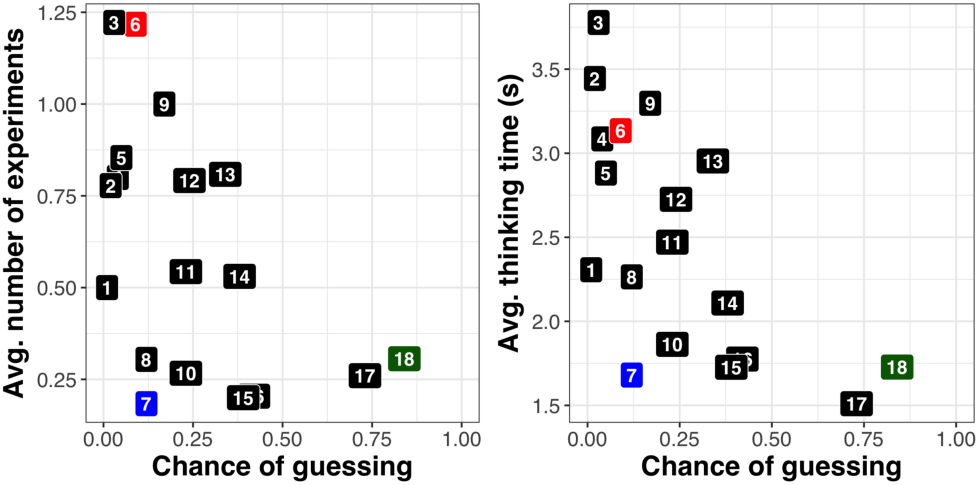
Plot of trial difficulty (chance of random shot being successful) versus number of practice shots (*left*) and average thinking time (*right*) for each trial. Points are labeled with trial indicators. See Fig. 1 for trials by number, and Fig. 3 for results from colored trials.

**Figure 3:**
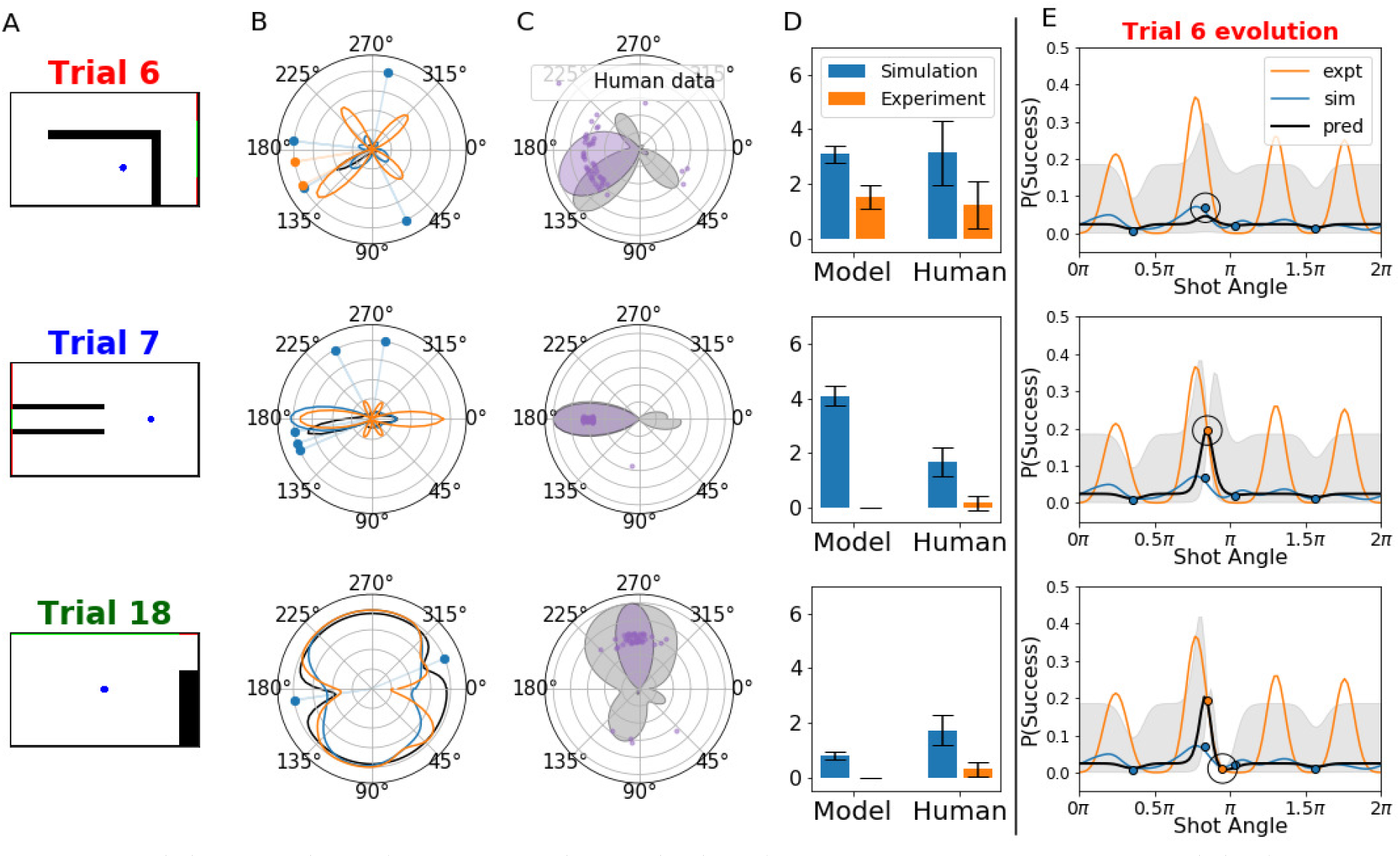
Qualitative behavior of the model. **A**: Screen shots of the three different trials (see main text for logic behind selection of trials). **B**: Polar plots showing the results of one run of the model, with probability of success from simulation and experiment. Dots indicate observations with blue dots marking simulations and orange dots marking experiments. The black line indicates the final estimate from the model (from that run) for probability of success. **C**: Density of final shots taken by the model, overlaid with final shots taken by participants. **D**: Number of experiments and simulations as generated by our model and humans (with amount of time spent not doing an experiments before taking the final shot as a proxy for number of simulations, with an assumption of 0.7 s/simulation, for ease of visualization) **E**: Evolution of one run of the model’s prediction over simulation/action choices, from top to bottom. The black line is the model’s belief (with grey indicating ± 1sd), the blue line is simulation probability, and the orange line is experimentation probability; dots indicate observations, with the same corresponding colors, and circles are the most recent observation. *Top:* the model initially simulates which increases probability of success in the region but is still uncertain. *Middle:* an experiment gives much more certainty, but it is still unclear if clockwise (right) or anticlockwise (left) from that point is better. *Bottom:* the final experiment updates belief that shooting clockwise from the first experiment would likely miss.

Participants were sensitive to trial features beyond this gross difficulty measure. Even for trials with similar probabilities of guessing, there was a wide variation in the average number of practice shots participants took; including trial indicators provided better predictions of the number of practice shots participants used as compared to predictions based only on the chance of a successful guess (*χ*^2^(16) = 115, *p* < .001). This can also be observed when comparing specific trials: for example, looking at the difference between Trial 7 (blue in Fig. 2), and both Trial 6 (red) and Trial 18 (green). While Trial 6 & 7 have roughly the same “chance” difficulty (9% vs 12%), participants rarely took practice shots in Trial 7 (0.18 / participant) but used a large number of practices for Trial 6 (1.21 / participant). Conversely, while the random chance of success are very different between Trial 7 & 18 (12% vs 84%) participants use a comparable number of practice shots in both (0.18 / participant for Trial 7 and 0.31 / participant for Trial 18).

We next investigated whether executing physical experiments improved participants’ performance across participants; however, we found no evidence for this. There was no evidence that participants who performed more practice shots were more accurate (*r*(48) = −0.07, *p* = .62), nor did they spend less time thinking (*r*(48) = −0.16, *p* = .26). While we would expect that experimentation would improve performance, it is possible that less skilled participants need this additional information, while more skilled participants can perform just as well without it.

In summary, participants modulated their simulation/experiment policy in response to trial difficulty, but not to relative action cost. Next, we turn to a model that can integrate information from simulation and physical actions to explain this pattern of information seeking across trials.

## Computational model

The integration model was designed to map actions (shot angles) to outcomes (probability of hitting the goal) by combining information from both simulation and experimentation. Let θ be the shooting angle and*f*(θ) be the log odds that θ will lead to a target hit. People learn this function by sampling physical paths, either through simulation or an experiment. Simulations are derived from the probability of hitting the target using the noisy physics model of Smith & Vul (2013), and experimental outcomes are sampled from a smoothed function mapping the actions to actual outcomes according to the physics engine used in the experiment.

To capture how people generalize from sampled paths to new paths, we assume that people learn a Gaussian process (GP) regression model of *f*(θ) (Rasmussen & Williams, 2006), a flexible nonparametric framework for function learning that has previously been successfully applied to model mental simulations (Hamrick & Griffiths, 2014). A GP prior over functions is specified by a mean (typically assumed to be0) and a covariance function, or kernel, *k* (θ,θ^′^) which governs the smoothness of the function. Given this prior, the posterior over functions is Gaussian and can be computed in closed form (Rasmussen & Williams, 2006).

While mental simulations are fast, they are also noisier than physical experiments, and hence not as informative. To integrate information from both of these sources of information, we follow the proposal put forward by Marco et al. (2017) for a kernel that can integrate simulations and experiments: 
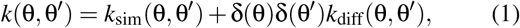
where δ(θ) = 1 indicates that θ was used to generate an experiment, and δ(θ) = 0 indicates that θ was used to generate a simulation. The kernel*k*_sim_ models the contribution from simulations, and*k*_diff_ models the difference between simulation and experiment. This is similar to exemplar-based models that generalize based on the distance (similarity) to previous observations (Griffiths et al., 2009), except observations here can be either simulations or experiments. To account for the differences in uncertainty from these two sources, we weight the distances to previous examples differently for each source, with a weaker generalization for simulations (using kernel *k*_sim_) and a stronger one for experiments (using kernel *k*_sim_ + *k*_diff_). We use a radial basis function for both kernels.

In order to find the angle θ* that optimizes *f*(θ), our model has to query informative angles. When choosing angles to query, it also has to account for a differences in costs of each information source. This requires us to have a principled way to quantify how valuable an observation from each of these sources of information is, relative to its cost. One such measure is how much an observation from each source reduces the entropy over the maxima of *f*(θ).

An acquisition function that utilizes this metric is predictive entropy search (PES, Hernández-Lobato et al., 2014). We adapt this acquisition function following Marco et al. (2017) to accommodate the two information sources. The mapping between query angle θ and how much it reduces entropy over the maxima of *f*(θ) is given by PES_sim_(θ) and PES_expt_(θ), depending on if the query is a simulation or a physical experiment. The simulations are more noisy so PES_sim_(θ) < PES_expt_(θ). To account for the difference in cost, we associate an effort measure with both kinds of evaluations *t*_sim_ for the simulation and *t*_exp_ for the physical experiments. While simulations are less informative, they are normally cheaper than experiments, so *t*_sim_ < *t*_exp_. We then select the next angle to evaluate, θ_*n*+1_, and whether it is a simulation or experiment, δ(θ_*n*+1_), according to argmax_θ,*i*∈{sim,exp}_ PES_*i*_(θ)/*t_i_*. The predictive entropy search first tries four randomly selected mental simulations, and afterwards chooses to evaluate new points based on the PES acquisition function.

We discretized the possible shots into 100 different angles spanning the circle, and precomputed for each trial and angle the following variables: (a) whether that shot would hit the green goal according to the game physics, and (b) the probability that a simulation would hit the green goal based on 50 random simulations from the noisy physics model. These variables are then smoothed using a Gaussian window, to account for perceptual error in distinguishing angles. We then take the logit of these functions to transform them onto the real line in order to carry out unconstrained Gaussian process regression.

There are four free parameters in this model: the ratio of the cost of simulations versus experiments (*t*_exp_/*t*_sim_), the simulation noise variance relative to the experiment noise variance, and two stopping criteria for the optimization: an entropy threshold that stops if a lower bound for the relative change in PES is reached, and a probability threshold if an upper bound for the best-so-far probability of success is reached. We used grid search to select the parameter values that minimized the sum-squared error between the number of experiments predicted by the model and the participants.*

## Model results

Our model closely tracked the frequency of experiments participants chose to perform across trials, *r*(16) = 0.68, *p* = 0.002 (Fig. 4, left). We then compared the number of simulations done by the model with the number of simulations done by participants. Since the number of simulations done by a participant is not explicitly available, a rough proxy is the amount of time participants spent thinking. We found a moderate correlation between the predicted number of simulations and participant’s thinking time (*r*(16) = 0.48, *p* = .042; Fig. 4, right).

**Figure 4:**
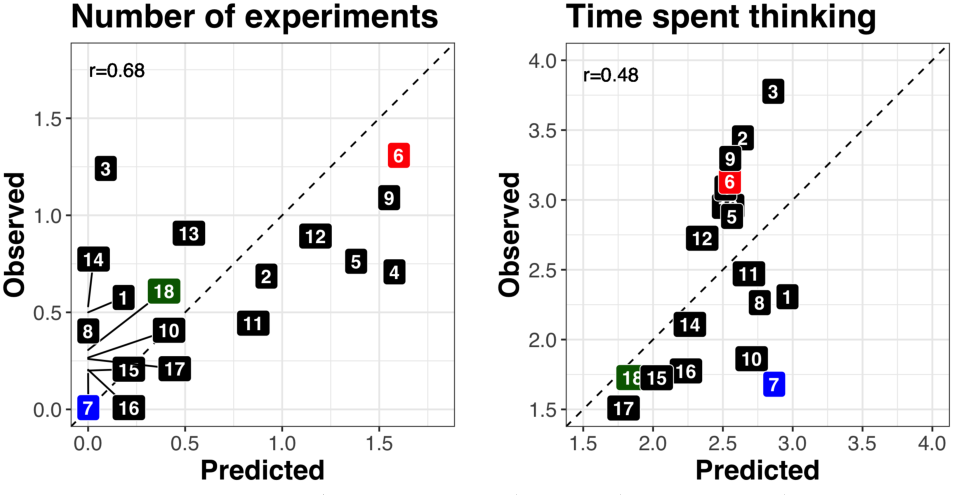
Correlations between model predictions and average participant behavior for each trial. *Left*: number of experiments. *Right*: time spent thinking (calculated based on a linear regression from number of model simulations, in seconds).

While there was a strong correlation in participants’ data between the number of practice shots and thinking time per trial (*r*(16) = 0.93, *p* < .001), the model predicts that there should be no appreciable correlation between the number of experiments and simulations (*r*(16) = .15, *p* = .54). Nonetheless, the number of simulations expected by the model does predict part of the unexplained variance in thinking time above and beyond the number of practice shots (*F*(1,15)= 4.55, *p* = .05), indicating that the number of simulations predicted by the model is still reflected in the amount of time spent thinking. Further, the correlation between simulations and thinking time (*r* = 0.48) is almost unchanged when partialing out the number of observed practice shots per trial (*r_part_* = 0.48), indicating that the model’s predicted simulations are explaining independent information to the number of practice shots.

We also see that our model is able to replicate the qualitative patterns of information seeking behavior in humans across trials, as highlighted by Trials 6, 7 and 18 in Figure 3: trials that are intuitively obvious and have very few practice shots (Trials 7 & 18) are solved by the model using simulation alone, while more difficult trials that spur participants to take practice shots (e.g., Trial 6) also spur the integration model to experiment.

In order to highlight the importance of integrating simulation and experiment for explaining the human data, we also test a model that cannot simulate, and fit it to the number of observed experiments from the data with a grid search through the stopping criteria parameters (the other parameters do not apply to this lesioned model). This model by definition cannot explain thinking time, though it can predict the number of practice shots (*r*(16) = 0.42, *p* = 0.08) which is not significantly worse than the full model (*z* = 0.21, *p* = 0.83). However, this is because the lesioned model was fit to the number of practice shots. Next, we investigate how the model uses those experiments to produce a final shot. Assessing the log-likelihood (LLH) of participants’ final shots under a smoothed distribution of model-predicted shots, we find that the full model (*LLH_full_* = −968, estimated kernel bandwidth = 0.295 radians) predicts participants’ shots better than the lesioned model (*LLH_lesion_*=−1,563, estimated bandwidth = 2.7 radians), making each final shot 2.02 times more likely to have occurred under the full vs. the lesioned model.

In summary, the lesioned, experiment-only model is not an adequate account of the data, and because we did observe practice shots, a simulation-only model could not account for participants’ behavior either. Our integrated model provides a good qualitative explanation for the information seeking behavior we see across trials and corresponds well to the final shots people take. It also provides reasonably good quantitative correlations with the number of experiments humans do across different trials, as well as the time they spent thinking.

## Discussion

Using a novel experimental paradigm that requires integrating simulation and experimentation to gather information, we found that people use both strategies, and furthermore that people’s use of both of these strategies varies across different scenarios. We can explain much of the variation in experimentation, time spent thinking, and final decisions using a single information gathering model that treats knowledge gained from both simulations and experiments in a common currency and uses an expected cost/benefit analysis to determine which strategy to use next.

This finding provides a starting framework to address the question of how people can flexibly integrate cheap but noisy information gathering in their heads and accurate but potentially expensive information gathering in the world. We find that people’s behavior is roughly consistent with a rational process model that chooses actions based on a measure of information gain. While this experiment focused on physical action planning, there are many domains in which people can gather information from both thinking and acting that this framework could be applied to, from problem solving to reinforcement learning (e.g., Gershman et al., 2017).

This is not the only possible model of people’s active information seeking (cf. Nelson, 2005): there are other ways of integrating information, assessing the expected benefits from each information source, or determining whether to simulate or act based on the current information. In future work we will assess further information seeking models that integrate simulation and experiment.

### Limitations

There are features of this integration problem that we will need to further investigate. For instance, according to our model there should be relatively little correlation between experiments and simulation, yet we find a large correlation between practice shots and thinking time in the empirical data. By assuming that thinking time is linearly related to the number of simulations, we could explain some of the variance in thinking time across trials, but left a large part unexplained. Future work needs to determine whether this is because our model of thinking time is incorrect (e.g., it should include time to set up practice shots), or if this points to an area of the integration process that requires further consideration.

Our participants were extremely consistent in the ultimate angle at which they shot the ball – in almost all cases choosing almost identical or one of two directions (Fig. 5, left). The integration model could explain why there were some accurate shots that people did not consider, but still picked out some angles that people did not (e.g., shooting at the opposite wall from a straight shot in Trials 7 & 18, Fig. 3). This may reflect preferences not built into the model (e.g., for shorter path lengths) or priors on the action-outcome mapping that can be gleaned before any simulations or actions occur.

**Figure 5:**
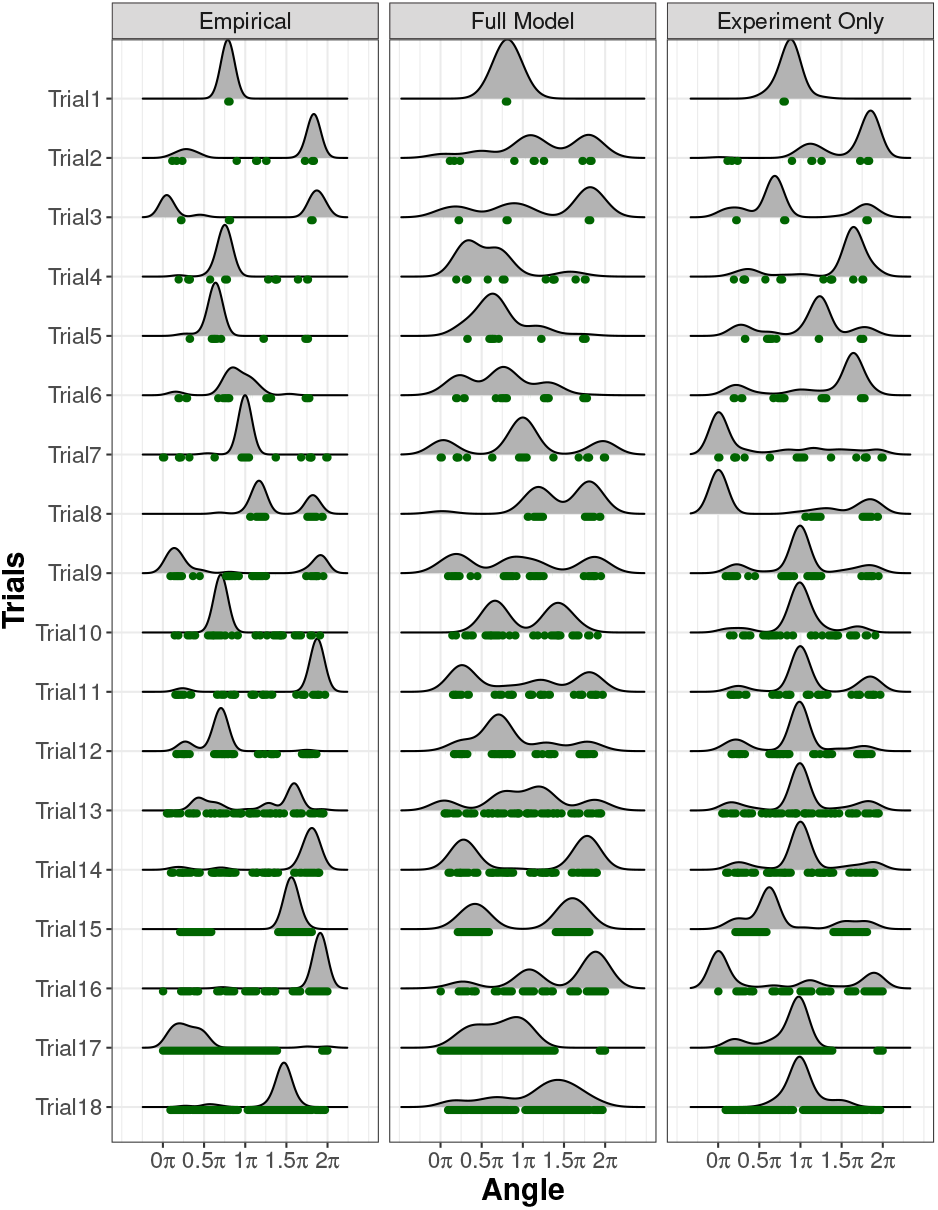
Density of final shots taken for each trial by participants (*left*), the full model (*middle*), and the experiment-only model (*right*). The green areas below the axes indicate angles that would be successful.

And finally, if we assume that people are sensitive to the costs of simulation versus action, we would expect to see a difference in behavior when experiments were cheaper or more costly, yet none was found here. Because this manipulation was performed across participants, perhaps people had set different expectations for points in each condition, and so any cost trade-offs would cancel out. This theory would be consistent with findings that between-subject incentive manipulations can be ineffective while the same incentive manipulations are effective within-subject (Harley, 1965).

### Conclusion

Mental simulation is a powerful tool that can help us plan ahead and solve complex problems without having to perform an action. Nonetheless, complex tasks often require both mental simulation and actual experiments. We have developed both a computational framework and reported preliminary behavioral findings that take a step towards understanding how people integrate simulation and experiments.

## Acknowledgments

ID is supported by Microsoft Research. KAS, SJG and JBT are supported by CBMM funded by NSF STC award CCF-1231216, and ONR grant N00014-13-1-0333. ES is supported by the Harvard Data Science Initiative. Code and data are available at: github.com/ishita-dg/SimulationVSAction

## Footnotes

* Our results are robust to fairly large variations in the parameter settings we searched. The only parameter that explained a large portion (32%) of the variance in the sum of squared errors over all parameter settings was the cost factor between experiment and simulation, with other factors explaining less than 1% of the variance.

